# *FMR1* loss results in early changes to intrinsic membrane excitability in human cellular models

**DOI:** 10.1101/2020.01.28.923425

**Authors:** Sara G. Susco, Mario A. Arias-Garcia, Violeta G. Lopez-Huerta, Amanda Beccard, Anne M. Bara, Jessica Moffitt, Justin Korn, Zhanyan Fu, Lindy E. Barrett

## Abstract

*Fragile X mental retardation 1 (FMR1)* encodes the RNA binding protein FMRP. Loss of FMRP drives Fragile X syndrome (FXS), the leading inherited cause of intellectual disability and a leading monogenic cause of autism. Cortical hyperexcitability is a hallmark of FXS, however, the underlying mechanisms reported, including alterations in synaptic transmission and ion channel expression and properties, are heterogeneous and at times contradictory. Here, we generated isogenic *FMR1*^*y*/+^ and *FMR1*^*y*/-^ human pluripotent stem cell (hPSC) lines using CRISPR-Cas9, differentiated these stem cell tools into excitatory cortical neurons and systematically assessed the impact of FMRP loss on intrinsic membrane and synaptic properties over the course of *in vitro* differentiation. Using whole-cell patch clamp analyses at five separate time-points, we observed significant changes in multiple metrics following FMRP loss, including decreased membrane resistance, increased capacitance, decreased action potential half-width and higher maximum frequency, consistent with *FMR1*^*y*/-^ neurons overall showing an increased intrinsic membrane excitability compared with age-matched *FMR1*^*y*/+^ controls. Surprisingly, a majority of these changes emerged early during *in vitro* differentiation and some were not stable over time. Although we detected significant differences in intrinsic properties, no discernable alterations were observed in synaptic transmission. Collectively, this study provides a new isogenic hPSC model to study the mechanisms of *FMR1* gene function, identifies electrophysiological impacts of FMRP loss on human excitatory cortical neurons over time *in vitro*, and underscores that early developmental changes to intrinsic membrane properties may be a critical cellular pathology contributing to cortical hyperexcitability in FXS.

## Introduction

Fragile X syndrome (FXS) is caused by a repeat expansion in the 5’UTR of *Fragile X mental retardation 1* (*FMR1*), leading to loss of Fragile X mental retardation protein (FMRP)(1–3). Studies using EEG and fMRI have revealed significant differences in cortical excitability between FXS patients and unaffected controls (4–6). Paralleling these results, a large body of experimental research in animal models reports altered excitability in the cerebral cortex following FMRP loss (7–9), which may be due to mis-regulation of RNA targets encoding synapse-associated proteins. While human pluripotent stem cell (hPSC) models of FXS hold great promise for dissection of human disease relevant mechanisms and drug screening, they have yielded diverse, and at times conflicting data on the impact of FMRP loss on human neurite development, synaptic connectivity and intrinsic and synaptic properties (7, 10–12). Even fundamental measurements such as neurite outgrowth have been reported to be decreased (10, 11), increased (12) or unchanged (7) following FMRP loss in different hPSC models. A combination of technical and experimental variables such as differences in genetic backgrounds, constitutive versus inducible FMRP loss, different cell type(s) and cell ratios generated by distinct *in vitro* differentiation paradigms, inclusion or exclusion of multiple additional cell types in co-cultures such as mouse neurons or glia, and different time-points of analyses all likely contribute to varying phenotypic presentations. This further implies that some FXS cellular phenotypes may be sufficiently subtle as to not rise above these technical and biological sources of variation (7), complicating cross study comparisons and extrapolation to the human disease state *in vivo*. While the lack of congruity across studies presents a challenge, careful assessment of different sources of variance will likely facilitate a more holistic view of the impact of FMRP loss on neuronal development, synaptic connectivity and function in the human brain. In particular, a majority of studies of the impacts of FMRP loss on physiological function focus on a single time-point of analysis, which led us to consider the initial pathological changes due to FMRP loss and how such changes dynamically evolve over different stages of differentiation.

Specifically, we utilized CRISPR-Cas9 technology to generate isogenic hPSC lines with or without constitutive FMRP loss and used these cellular tools to generate excitatory cortical neurons from a well-established *in vitro* differentiation paradigm (7, 13–22). To gain insight into the physiological response to *FMR1* loss over the course of *in vitro* differentiation, we measured intrinsic membrane properties and synaptic transmission in isogenic *FMR1*^*y*/+^ and *FMR1*^*y*/-^ neurons at five separate time-points from two to five weeks of *in vitro* differentiation. We identified significant cell intrinsic defects but not synaptic transmission deficits driven by FMRP loss in these cellular systems. Moreover, these differences began to emerge relatively early during *in vitro* differentiation and several metrics dynamically changed over time, which may account for some of the previous discrepancies reported in the literature.

## Results

To facilitate analyses of FMRP loss isolated from differences in genetic background, we first generated and validated *FMR1*^*y*/+^ and *FMR1*^*y*/-^ isogenic hPSC lines (referred to as *FMR1* WT and *FMR1* KO, respectively) by targeting exon 4 of *FMR1* with CRISPR-Cas9 in the XY line H1 (**Fig. 1A**). Indels in exon 4 of *FMR1* led to complete loss of FMRP expression (**Fig. 1B**) and loss of FMRP did not prevent hPSC differentiation into excitatory neurons (**Fig. 1C**), consistent with previous reports using a similar differentiation paradigm (7). Targeted cell lines maintained pluripotency, tri-lineage potential and karyotype stability (**Fig. S1 and data not shown**).

**Fig. 1.**
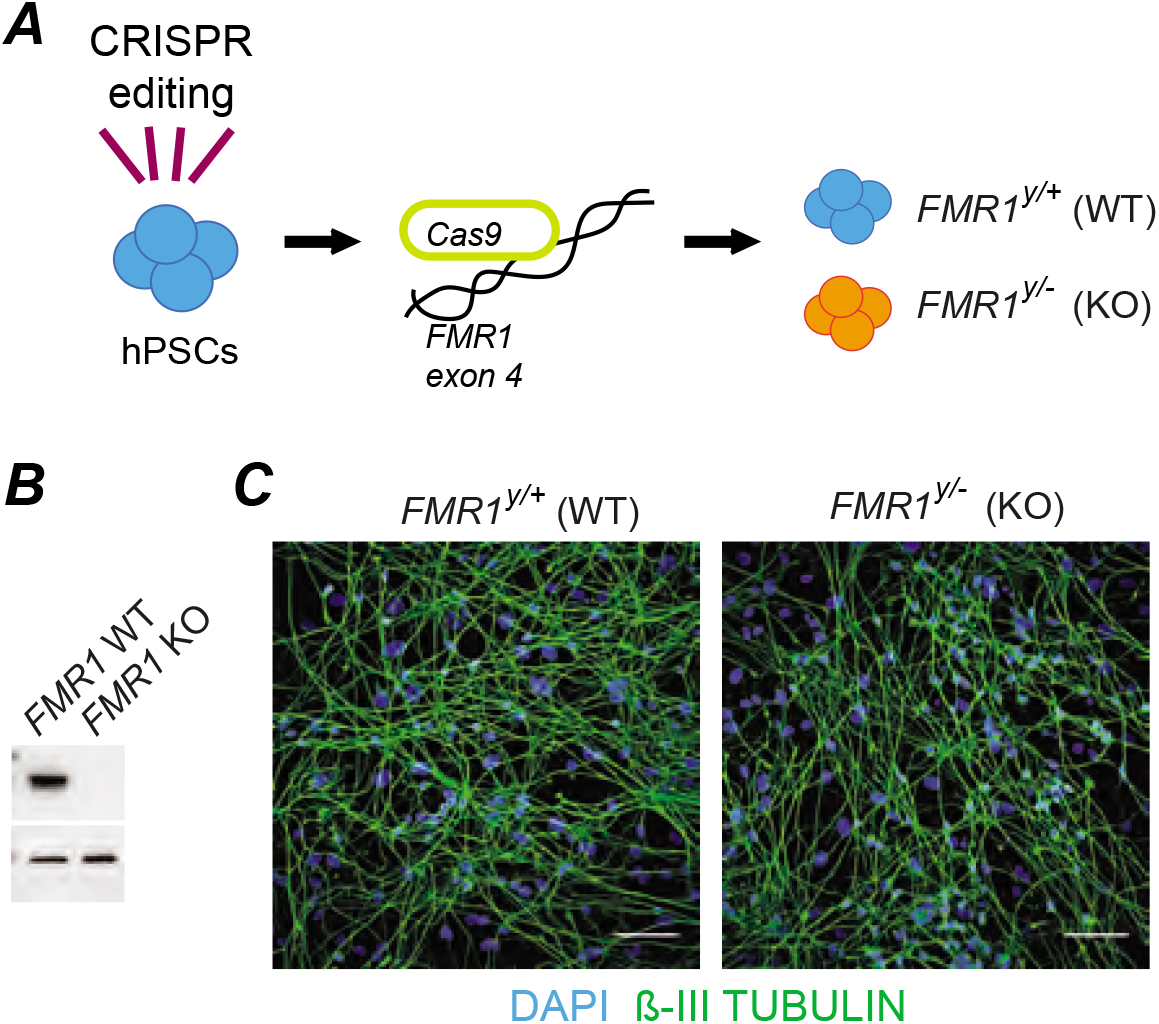
Generation of isogenic *FMR1* cellular resources. **A.** Schematic of CRISPR-Cas9 editing strategy targeting exon 4 of *FMR1* to generate isogenic *FMR1*^*y*/+^ (WT) and *FMR1*^*y*/-^ (KO) hPSC lines. **B.** Western blot analysis showing loss of FMRP in CRISPR edited *FMR1* KO hPSCs compared to *FMR1* WT hPSCs. **C.** Expression of β-III-Tubulin (green) in *FMR1* WT (left) and *FMR1* KO (right) neurons. Cells are counterstained with DAPI (blue). Scale bar = 50μm.

To understand how constitutive loss of FMRP affects the electrophysiological profiles of human excitatory neurons, we first assessed intrinsic membrane properties. For all whole-cell patch clamp experiments, *FMR1* WT and *FMR1* KO neurons were plated at densities of 40,000 cells/cm^2^ with mouse glia on glass coverslips. hPSCs used for *in vitro* differentiation were infected with CAMK2A-GFP or CAMK2A-mOrange and only GFP+ or mOrange+ neurons were used for patch clamp analyses in order to select for neurons sufficiently mature as to express CAMK2A, as previously described (13). Neurons from each genotype were recorded from up to five separate batches of *in vitro* differentiation at five separate time-points from days 11 to 35 (D11, D14, D21, D28 and D35) (**Fig. S2**). As shown in **Fig. 2A**, we first assessed passive membrane properties of the neurons including: resting membrane potential (RMP), membrane resistance (Rm), membrane time constant (Tau) and membrane capacitance (Cm). From these properties, we observed significant differences in Rm and Cm in *FMR1* KO compared to *FMR1* WT neurons, but no changes to RMP or Tau at any time-point measured (**Fig. 2B-E**). Specifically, we observed decreased Rm and increased Cm in *FMR1* KO neurons relative to *FMR1* WT neurons (**Figs. 2C, E**). However, only changes to Rm remained significant across time from D14-D35 (**Fig. 2C**), while Cm appeared dynamic, with significant differences observed at D14 and D35 and no significant differences at D11, D21 or D28 (**Fig. 2E**). While both significant parameters (i.e., decreased Rm and increased Cm in *FMR1* KO neurons relative to *FMR1* WT neurons) are consistent with *FMR1* KO neurons showing an increased maturation profile compared with age-matched *FMR1* WT control neurons, they also suggest that the neurons undergo dynamic shifts in their physiological properties over time in response to constitutive *FMR1* loss. Our data are also consistent with FMRP driving early (D14) changes to intrinsic membrane properties.

**Fig. 2.**
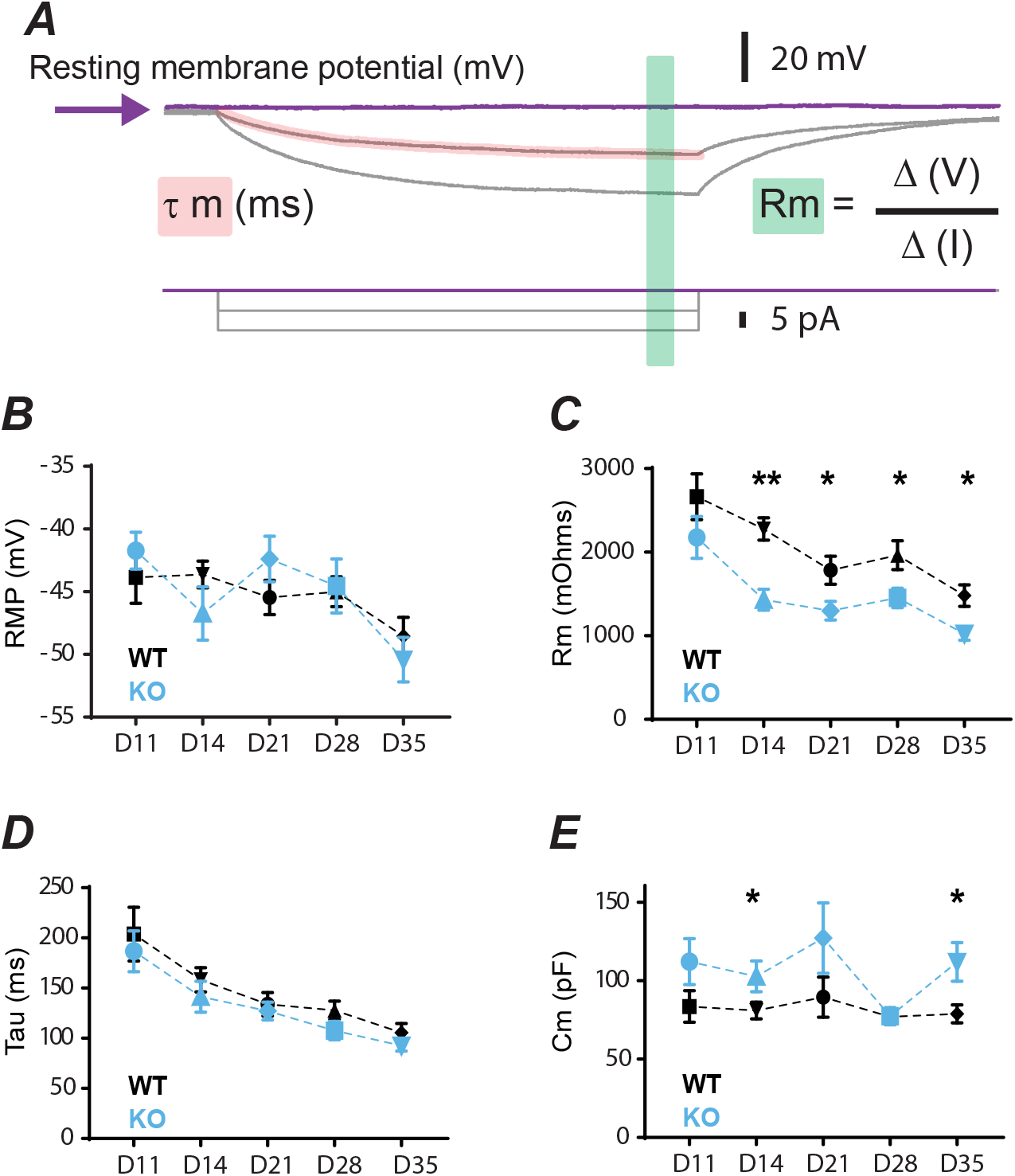
Constitutive *FMR1* loss leads to altered intrinsic membrane properties over time. **A.** Schematic of analysis parameters. **B.** Summary of resting membrane potential (RMP) measurements in *FMR1* WT (black) and *FMR1* KO (blue) neurons at indicated time-points. No significant differences were identified. **C.** Summary of membrane resistance (Rm) measurements in WT (black) and KO (blue) neurons at indicated time-points. KO neurons showed significantly lower membrane resistance than their WT counterparts from D14 onward (**p<0.01, *p<0.05, unpaired t-test) indicative of a higher level of maturation. **D.** Summary of membrane time constant (Tau) measurements in WT (black) and KO (blue) neurons at indicated time-points. No significant differences were identified. **E.** Summary of membrane capacitance (Cm) measurements in WT (black) and KO (blue) neurons at indicated time-points. KO neurons showed significantly higher Cm compared with their WT counterparts at D14 and D35 but not at D11, D21 or D28 (*p<0.05, unpaired t-test).

We next assessed the active membrane properties of the neurons, including action potential threshold (APthres), action potential half-width (APhw), action potential amplitude (APamp), after hyperpolarization (AHP) and maximum frequency (**Fig. 3A**). From these properties, we observed a significant increase in firing frequency at D21, D28 and D35 and decreased APhw at D11 and D14 in *FMR1* KO neurons relative to *FMR1* WT neurons, with no changes to APthres, APamp or AHP at any time-point measured (**Fig. 3B-G**). Significant differences in APhw were evident early during *in vitro* differentiation at D11 and D14 but these differences disappeared at later time-points (**Fig. 3B**), suggesting that neurons may undergoing compensatory changes over time. Significant differences in firing frequency curves and maximum frequency emerged after D21 and remained until D35 (**Fig. 3F-G**). As with decreased Rm and increased Cm in *FMR1* KO neurons relative to *FMR1* WT neurons described above (**Fig. 2C, E**), increased maximum frequency and decreased APhw are consistent with *FMR1* KO neurons showing an increased maturation profile compared with age-matched *FMR1* WT control neurons.

**Fig. 3.**
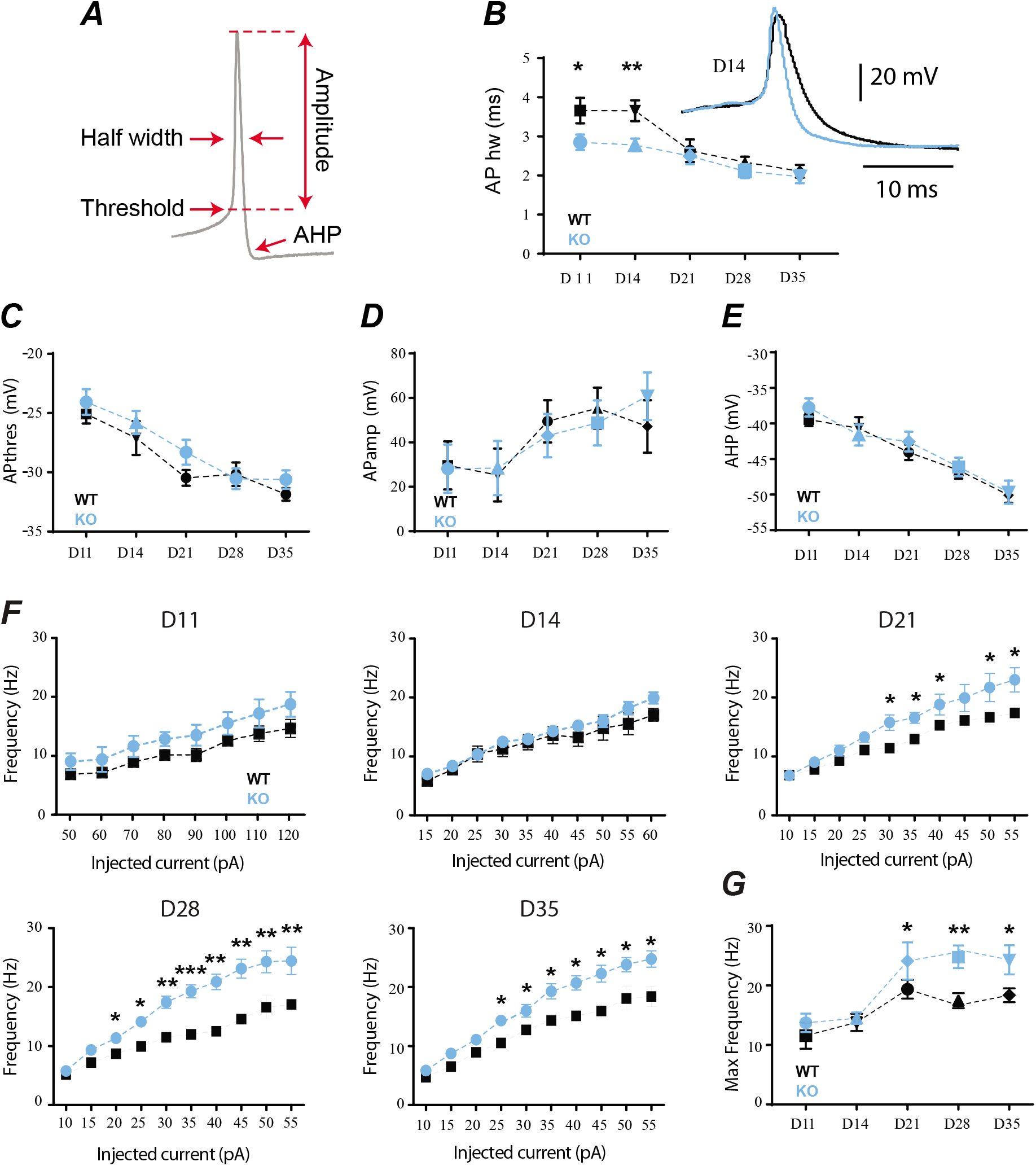
Constitutive *FMR1* loss leads to altered action potential properties over time. **A.** Schematic of action potential properties measured by whole-cell patch clamp analyses from five independent batches of *in vitro* differentiation across five time-points. **B.** Summary of AP half width measurements in *FMR1* WT (black) and *FMR1* KO (blue) neurons at indicated time-points. KO neurons showed significantly shorter APhw compared to their WT counterparts at D11 and D14 (*p<0.05, **p<0.01, unpaired t-test). Inset shows a representative expanded action potential for WT and KO neurons at D14. **C-E.** Summary of APthres, APamp and AHP measurements in WT (black) and KO (blue) neurons at indicated time-points. No significant differences were identified. **F.** Summary of the firing frequency curve as a response to increasing current injection in WT (black) and KO (blue) neurons at indicated time-points. KO neurons showed significantly higher maximum frequencies compared with WT neurons from D21 onward (*p<0.05, **p<0.01, ***p<0.001, unpaired t-test). **G.** Summary of maximum frequency measurements in WT (black) and KO (blue) neurons at indicated time-points. KO neurons showed significantly higher maximum frequencies compared with WT neurons from D21 onward (*p<0.05, **p<0.01, unpaired t-test).

Finally, we measured synaptic transmission including mean amplitude and frequency of synaptic events as well as the percentage of neurons with spontaneous excitatory post-synaptic currents (sEPSCs) in the absence of the AP blocker tetrodotoxin (TTX) **(Fig. 4)** from the same neurons used to assess intrinsic properties (**Figs. 2–3**). We did not detect differences in any of these synaptic properties measured under steady state conditions (**Fig. 4A-D**).

**Fig. 4.**
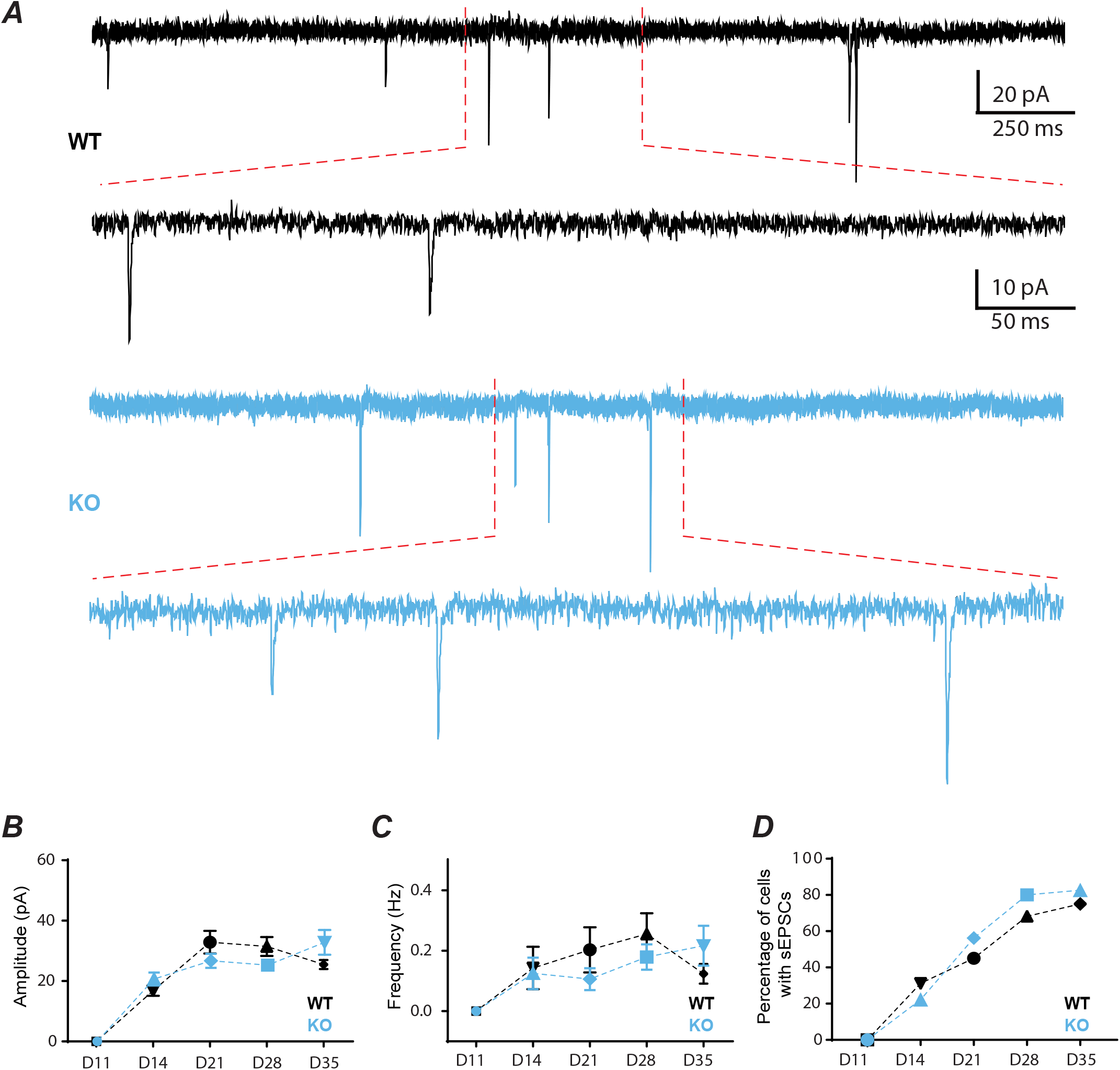
Constitutive *FMR1* loss does not lead to significant differences in synaptic properties at steady-state. **A.** Sample traces showing spontaneous excitatory postsynaptic currents (sEPSCs) in WT (black) and KO (blue) neurons at D35. **B-D.** Summary of sEPSC amplitude **(B)**, frequency **(C)** and percentage of cells that showed responses **(D)** in WT (black) and KO (blue) neurons at indicated time-points. No significant differences were identified under steady state conditions.

## Discussion

By generating isogenic *FMR1* WT and *FMR1* KO cell lines plated under identical culture conditions and recorded from up to five separate batches of *in vitro* differentiation over five separate time-points, we observed subtle but significant differences in intrinsic membrane properties that collectively suggest an increased physiological maturation profile of *FMR1* KO neurons compared to *FMR1* WT controls, consistent with previous studies describing electrophysiological and neural network activity in mouse models of FXS (23–28). Our results suggest that early developmental changes in the intrinsic membrane properties could potentially underlie subsequent cortical hyperexcitability and other related neurological deficits associated with FXS. Given that human *in vitro* derived neurons most closely resemble prenatal cell stages *in vivo* (13), these analyses underscore the importance for future studies to identify the ionic cellular mechanisms underlying the hyperexcitability in FXS during prenatal development. Of note, not all phenotypes were stable over time, and indeed we observed dynamic changes in multiple metrics, suggesting that neurons respond to constitutive FMRP loss as they differentiate *in vitro* and some discrepancies in the literature may be due to differing time-points of analysis. Indeed, a majority of studies using *in vitro* human neuronal of disease assess electrophysiological properties at a single time-point (7, 15, 16). Although we detected significant differences in intrinsic properties, we did not uncover alterations in synaptic function consistent with a previous study of FXS using the same neuronal cell type (7). This suggests that additional cell types or conditions may be required to unmask synaptic phenotypes *in vitro*. For example, our inclusion of mouse glia increased the health and viability of our neuronal cultures, but could also reduce phenotypic severity if glia contribute to electrophysiological phenotypes in FXS, as reported in other experimental contexts (29–31). While our experiments allow us to sensitively reveal phenotypes specific to excitatory cortical neurons, they cannot capture the complex relationship with inhibitory neurons and how this may drive network perturbations. Though our isogenic system eliminates variability in genetic background between patient and control samples and thus allows for the detection of more subtle phenotypes, given the variability reported in both FXS molecular phenotypes (e.g., differing levels of protein synthesis changes across patients)(32) as well as FXS clinical presentations (e.g., the fact that a subset of patients have seizures and a subset do not)(33), physiological phenotypes *in vitro* may similarly vary across different parental cell lines. Our study identifies specific time-points and conditions in human *in vitro* derived cortical neurons for further investigation. Future studies will be required to fully understand how different patient brain cell types respond to *FMR1* loss over time and how this impacts the development of more complex networks of human excitatory neurons, inhibitory neurons and astrocytes. hPSC models are well positioned to address increasingly complex questions related to the impact of a defined genetic change on multiple or mixed brain cell types to uncover key developmental mechanisms with relevance to human disease.

## Materials and Methods

### Stem cell culture and CRISPR-Cas9 based genome engineering

The human embryonic stem cell line H1 (34) was obtained commercially from WiCell Research Institute (https://www.wicell.org). All studies using H1 followed institutional IRB and ESCRO guidelines approved by Harvard University. Stem cell culture and assessment of pluripotency and tri-lineage potential was carried out as previously described (35–37). In brief, stem cells were grown and maintained in mTeSR1 medium (Stem Cell Technologies) on geltrex-coated plates (Life Technologies). Cells were routinely tested to confirm the absence of mycoplasma contamination (Lonza MycoAlert). CRISPR-Cas9 based genome engineering experiments were carried out as previously described (35–37). In brief, H1 was transfected with pre-assembled Cas9 protein (NEB) + crRNA and tracrRNA (Synthego), targeting full-length *FMR1* upstream of predicted functional domains (gRNA: GTTGGTGGTTAGCTAAAGTG), using the NEON system (Life Technologies). Wild-type cells are those that went through the gene-targeting protocol but were not edited. Antibodies used to assess pluripotency were anti-OCT3/4 (R&D Systems AF1759), anti-SOX2 (BD 245619), and anti-TRA-1-60 (Santa Cruz Biotechnology sc-21705). Antibodies used to assess tri-lineage potential were anti-AFP (Sigma A8452), anti-SMA (Sigma A2547), and anti-β-III-Tubulin (R&D Systems MAB1195).

### Generation of human excitatory cortical neurons

Human neurons were generated as previously described (13, 14). In brief, H1 hPSCs were transduced with TetO-Ngn2-T2A-Puro and Ubiq-rtTA lentivirus and treated with doxycycline to induce ectopic Ngn2 expression combined with the extrinsic addition of SMAD inhibitors (SB431542, 1614, Tocris, and LDN-193189, 04-0074, Stemgent), Wnt inhibitors (XAV939, 04-00046, Stemgent) and neurotrophins (BDNF, GDNF, CNTF) followed by puromycin treatment to eliminate uninfected stem cells. Ultra-high lentiviral titer was generated by Alstem, LLC.

### Whole-cell patch clamp analysis

Stem cells were infected with CAMK2A-GFP or CAMK2A-mOrange ultra-high titer lentivirus (Alstem, LLC) and differentiated into neurons as described above. At D4, neurons were plated at a density of 40,000 cells/cm^2^ on a bed of mouse glia on poly-D-lysine and laminin coated glass coverslips with Geltrex coating. Neurons were kept in NBM supplemented with B27 and neurotrophins. Whole-cell patch clamp recordings were performed at five time-points: Days 11, 14, 21, 28 and 35. Cultured neurons were transferred to a recording chamber and constantly perfused at a speed of 3ml/min with an extracellular solution containing (in mM): 119 NaCl, 2.3 KCl, 2 CaCl_2_, 1 MgCl_2_, 15 HEPES, 5 glucose, phenol red (0.25mg/L) and D-serine (10μM) (all from Sigma) adjusted to pH 7.2-7.4 with NaOH. Osmolarity was adjusted to 325 mOsm with sucrose. Recording pipettes (KG33, King Precision Glass) were pulled in a horizontal pipette puller (P-97, Sutter Instruments) with a tip resistance of 3–5 MΩ. Pipettes were filled with an internal solution containing in mM: 120 K-Gluconate, 2 MgCl_2_, 10 HEPES, 0.5 EGTA, 0.2 Na2ATP, and 0.2 Na_3_GTP. All experiments were performed at room temperature. The recordings were made with a microelectrode amplifier with bridge and voltage clamp modes of operation (Multiclamp 700B, Molecular Devices). Cell membrane potential was held at −60 mV, unless specified otherwise. Signals were low-pass filtered at 2 kHz and sampled at 10kHz with a Digidata 1440A (Molecular Devices). All data were stored on a computer for subsequent off-line analysis. Cells in which the series resistance (Rs, typically 8–12 MΩ) changed by >20% were excluded for data analysis. In addition, cells with Rs more than 25 MΩ at any time during the recordings were discarded. Conventional characterization of neurons was made in voltage and current clamp configurations. The membrane resistance, time constant (tau) and capacitance were measured in current clamp mode as described previously (38). CAMK2A expressing neurons were identified for recordings on the basis of GFP or mOrange expression visualized with a microscope equipped with GFP and Texas red filter (BX-51WI, Olympus). The electrophysiologists were blinded to genotype until data analyses were complete. The number of individual neurons recorded from and the number of independent batches of neurons are shown in **Fig. S2.**

## Acknowledgements

We thank members of the Barrett and Fu labs for insightful discussions and critical reading of the manuscript and John Sherwood for statistical advice and insight.

---

**Fig. S1:**
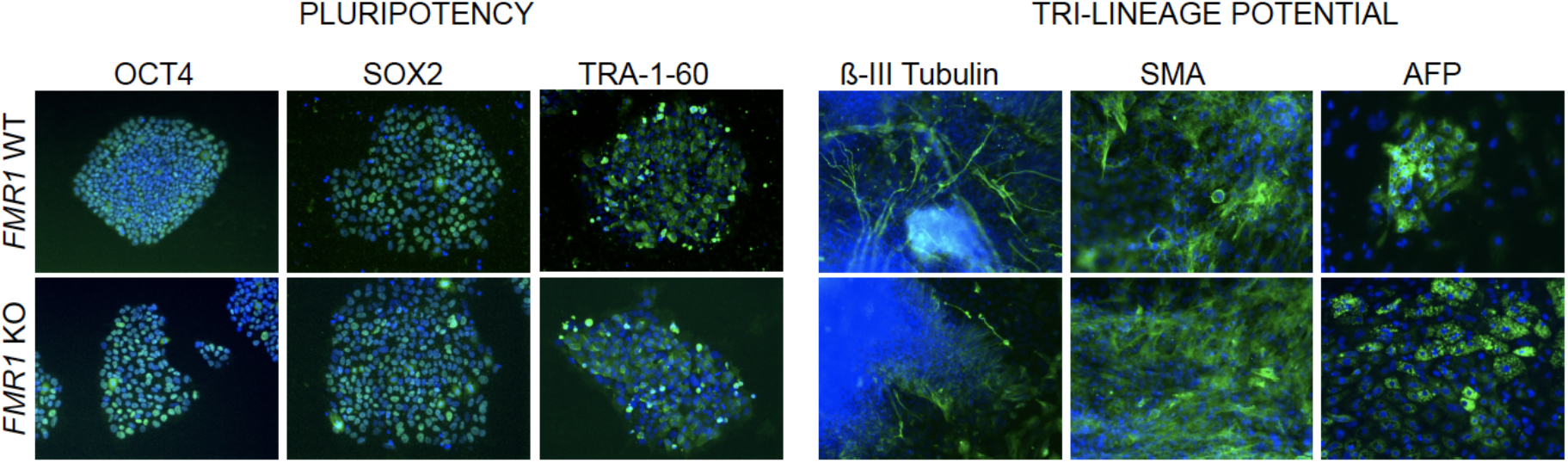
Assessment of pluripotency (left) and tri-lineage potential (right) for *FMR1* WT (top) and *FMR1* KO (bottom) hPSC lines.

**Fig. S2:**
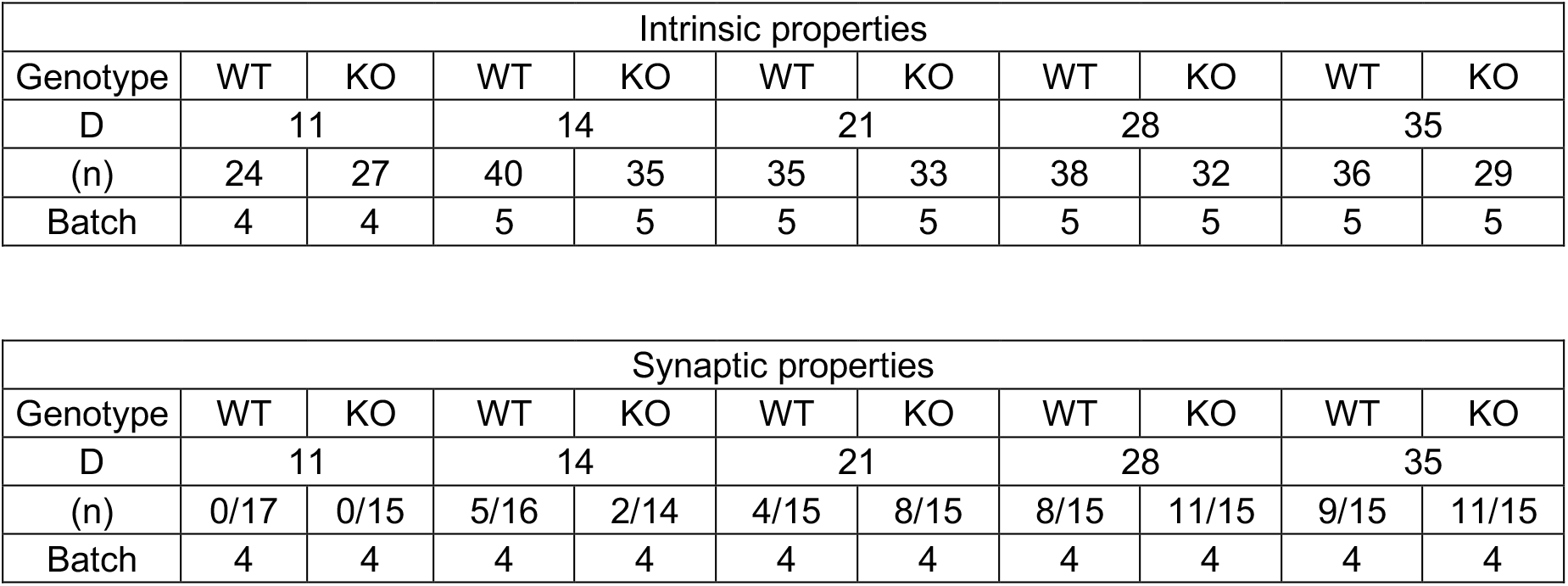
Chart showing the numbers of neurons and numbers of batches used for recordings. D=Days *in vitro*; (n)= number of neurons recorded from in total for each condition. Batch indicates the total number of independently generated sets of neurons utilized to measure the indicated properties.

## Other Information

### Author contributions

L.E.B., Z.F. and S.G.S. conceived the project and wrote the manuscript, S.G.S. performed stem cell and neuron culture. M.A.A-G. and V.G.L-H. performed and analyzed whole cell patch clamp experiments. A.B. and A.M.B. generated gene edited cell lines. J.M. and J.K. provided analytical support. and L.E.B. and Z.F. supervised the study and secured funding. All authors discussed results and edited the manuscript.

## Data availability

All data is available in the manuscript or the supplementary materials. Cell lines are available upon request with appropriate institutional approvals and following WiCell regulations for cell line distribution.

## References

1. Darnell JC, Van Driesche SJ, Zhang C, Hung KY, Mele A, Fraser CE, et al. FMRP stalls ribosomal translocation on mRNAs linked to synaptic function and autism. Cell. 2011;146(2):247–61.

2. Dictenberg JB, Swanger SA, Antar LN, Singer RH, Bassell GJ. A direct role for FMRP in activity-dependent dendritic mRNA transport links filopodial-spine morphogenesis to fragile X syndrome. Developmental cell. 2008;14(6):926–39.

3. Bagni C, Tassone F, Neri G, Hagerman R. Fragile X syndrome: causes, diagnosis, mechanisms, and therapeutics. The Journal of clinical investigation. 2012;122(12):4314–22.

4. Ethridge LE, White SP, Mosconi MW, Wang J, Pedapati EV, Erickson CA, et al. Neural synchronization deficits linked to cortical hyper-excitability and auditory hypersensitivity in fragile X syndrome. Mol Autism. 2017;8:22.

5. Berry-Kravis E. Epilepsy in fragile X syndrome. Dev Med Child Neurol. 2002;44(11):724–8.

6. Musumeci SA, Hagerman RJ, Ferri R, Bosco P, Dalla Bernardina B, Tassinari CA, et al. Epilepsy and EEG findings in males with fragile X syndrome. Epilepsia. 1999;40(8):1092–9.

7. Zhang Z, Marro SG, Zhang Y, Arendt KL, Patzke C, Zhou B, et al. The fragile X mutation impairs homeostatic plasticity in human neurons by blocking synaptic retinoic acid signaling. Science translational medicine. 2018;10(452).

8. Gibson JR, Bartley AF, Hays SA, Huber KM. Imbalance of neocortical excitation and inhibition and altered UP states reflect network hyperexcitability in the mouse model of fragile X syndrome. J Neurophysiol. 2008;100(5):2615–26.

9. Larson J, Jessen RE, Kim D, Fine AK, du Hoffmann J. Age-dependent and selective impairment of long-term potentiation in the anterior piriform cortex of mice lacking the fragile X mental retardation protein. The Journal of neuroscience: the official journal of the Society for Neuroscience. 2005;25(41):9460–9.

10. Doers ME, Musser MT, Nichol R, Berndt ER, Baker M, Gomez TM, et al. iPSC-derived forebrain neurons from FXS individuals show defects in initial neurite outgrowth. Stem Cells Dev. 2014;23(15):1777–87.

11. Sheridan SD, Theriault KM, Reis SA, Zhou F, Madison JM, Daheron L, et al. Epigenetic characterization of the FMR1 gene and aberrant neurodevelopment in human induced pluripotent stem cell models of fragile X syndrome. PloS one. 2011;6(10):e26203.

12. Boland MJ, Nazor KL, Tran HT, Szucs A, Lynch CL, Paredes R, et al. Molecular analyses of neurogenic defects in a human pluripotent stem cell model of fragile X syndrome. Brain. 2017;140(3):582–98.

13. Nehme R, Zuccaro E, Ghosh S, Li C, Sherwood J, Pietilainen O, et al. Combining NGN2 Programming with Developmental Patterning Generates Human Excitatory Neurons with NMDAR-Mediated Synaptic Transmission. Cell reports. 2018;23(8):2509–23.

14. Zhang Y, Pak C, Han Y, Ahlenius H, Zhang Z, Chanda S, et al. Rapid single-step induction of functional neurons from human pluripotent stem cells. Neuron. 2013;78(5):785–98.

15. Pak C, Danko T, Zhang Y, Aoto J, Anderson G, Maxeiner S, et al. Human Neuropsychiatric Disease Modeling using Conditional Deletion Reveals Synaptic Transmission Defects Caused by Heterozygous Mutations in NRXN1. Cell stem cell. 2015;17(3):316–28.

16. Yi F, Danko T, Botelho SC, Patzke C, Pak C, Wernig M, et al. Autism-associated SHANK3 haploinsufficiency causes Ih channelopathy in human neurons. Science. 2016;352(6286):aaf2669.

17. Lin YT, Seo J, Gao F, Feldman HM, Wen HL, Penney J, et al. APOE4 Causes Widespread Molecular and Cellular Alterations Associated with Alzheimer’s Disease Phenotypes in Human iPSC-Derived Brain Cell Types. Neuron. 2018;98(6):1141–54 e7.

18. Marro SG, Chanda S, Yang N, Janas JA, Valperga G, Trotter J, et al. Neuroligin-4 Regulates Excitatory Synaptic Transmission in Human Neurons. Neuron. 2019;103(4):617–26 e6.

19. Meijer M, Rehbach K, Brunner JW, Classen JA, Lammertse HCA, van Linge LA, et al. A Single-Cell Model for Synaptic Transmission and Plasticity in Human iPSC-Derived Neurons. Cell reports. 2019;27(7):2199–211 e6.

20. Busskamp V, Lewis NE, Guye P, Ng AH, Shipman SL, Byrne SM, et al. Rapid neurogenesis through transcriptional activation in human stem cells. Molecular systems biology. 2014;10:760.

21. Deneault E, White SH, Rodrigues DC, Ross PJ, Faheem M, Zaslavsky K, et al. Complete Disruption of Autism-Susceptibility Genes by Gene Editing Predominantly Reduces Functional Connectivity of Isogenic Human Neurons. Stem cell reports. 2018;11(5):1211–25.

22. Tian R, Gachechiladze MA, Ludwig CH, Laurie MT, Hong JY, Nathaniel D, et al. CRISPR Interference-Based Platform for Multimodal Genetic Screens in Human iPSC-Derived Neurons. Neuron. 2019;104(2):239–55 e12.

23. Lovelace JW, Wen TH, Reinhard S, Hsu MS, Sidhu H, Ethell IM, et al. Matrix metalloproteinase-9 deletion rescues auditory evoked potential habituation deficit in a mouse model of Fragile X Syndrome. Neurobiology of disease. 2016;89:126–35.

24. Goncalves JT, Anstey JE, Golshani P, Portera-Cailliau C. Circuit level defects in the developing neocortex of Fragile X mice. Nature neuroscience. 2013;16(7):903–9.

25. Truszkowski TL, James EJ, Hasan M, Wishard TJ, Liu Z, Pratt KG, et al. Fragile X mental retardation protein knockdown in the developing Xenopus tadpole optic tectum results in enhanced feedforward inhibition and behavioral deficits. Neural Dev. 2016;11(1):14.

26. Schaefer TL, Davenport MH, Grainger LM, Robinson CK, Earnheart AT, Stegman MS, et al. Acamprosate in a mouse model of fragile X syndrome: modulation of spontaneous cortical activity, ERK1/2 activation, locomotor behavior, and anxiety. Journal of neurodevelopmental disorders. 2017;9:6.

27. Danesi C, Achuta VS, Corcoran P, Peteri UK, Turconi G, Matsui N, et al. Increased Calcium Influx through L-type Calcium Channels in Human and Mouse Neural Progenitors Lacking Fragile X Mental Retardation Protein. Stem cell reports. 2018;11(6):1449–61.

28. Lovelace JW, Rais M, Palacios AR, Shuai XS, Bishay S, Popa O, et al. Deletion of Fmr1 from Forebrain Excitatory Neurons Triggers Abnormal Cellular, EEG, and Behavioral Phenotypes in the Auditory Cortex of a Mouse Model of Fragile X Syndrome. Cerebral cortex. 2019.

29. Higashimori H, Schin CS, Chiang MS, Morel L, Shoneye TA, Nelson DL, et al. Selective Deletion of Astroglial FMRP Dysregulates Glutamate Transporter GLT1 and Contributes to Fragile X Syndrome Phenotypes In Vivo. The Journal of neuroscience: the official journal of the Society for Neuroscience. 2016;36(27):7079–94.

30. Jawaid S, Kidd GJ, Wang J, Swetlik C, Dutta R, Trapp BD. Alterations in CA1 hippocampal synapses in a mouse model of fragile X syndrome. Glia. 2018;66(4):789–800.

31. Jacobs S, Doering LC. Astrocytes prevent abnormal neuronal development in the fragile x mouse. The Journal of neuroscience: the official journal of the Society for Neuroscience. 2010;30(12):4508–14.

32. Jacquemont S, Pacini L, Jonch AE, Cencelli G, Rozenberg I, He Y, et al. Protein synthesis levels are increased in a subset of individuals with fragile X syndrome. Human molecular genetics. 2018;27(21):3825.

33. Kaufmann WE, Kidd SA, Andrews HF, Budimirovic DB, Esler A, Haas-Givler B, et al. Autism Spectrum Disorder in Fragile X Syndrome: Cooccurring Conditions and Current Treatment. Pediatrics. 2017;139(Suppl 3):S194–S206.

34. Thomson JA, Itskovitz-Eldor J, Shapiro SS, Waknitz MA, Swiergiel JJ, Marshall VS, et al. Embryonic stem cell lines derived from human blastocysts. Science. 1998;282(5391):1145–7.

35. Hazelbaker DZ, Beccard A, Bara AM, Dabkowski N, Messana A, Mazzucato P, et al. A Scaled Framework for CRISPR Editing of Human Pluripotent Stem Cells to Study Psychiatric Disease. Stem cell reports. 2017;9(4):1315–27.

36. Bara AM, Messana A, Herring A, Hazelbaker DZ, Eggan K, Barrett LE. Generation of a TLE3 heterozygous knockout human embryonic stem cell line using CRISPR-Cas9. Stem Cell Res. 2016;17(2):441–3.

37. Herring A, Messana A, Bara AM, Hazelbaker DZ, Eggan K, Barrett LE. Generation of a TLE1 homozygous knockout human embryonic stem cell line using CRISPR-Cas9. Stem Cell Res. 2016;17(2):430–2.

38. Golowasch J, Thomas G, Taylor AL, Patel A, Pineda A, Khalil C, et al. Membrane capacitance measurements revisited: dependence of capacitance value on measurement method in nonisopotential neurons. Journal of Neurophysiology. 2009;102(4):2161–75.

